# Fast MAS NMR Spectroscopy Can Identify G-Quartets and Double-Stranded Structures in Aggregates Formed by GGGGCC RNA Repeats

**DOI:** 10.1101/2025.09.07.674584

**Authors:** Sara Zager, Nataša Medved, Mirko Cevec, Urša Čerček, Boris Rogelj, Janez Plavec, Jaka Kragelj

## Abstract

The expansion of GGGGCC repeats within the *C9orf72* gene has been linked to amyotrophic lateral sclerosis (ALS) and frontotemporal dementia (FTD). In neurons from patients with expanded repeats, *C9orf72* GGGGCC repeat RNA predominantly forms nuclear foci, and *in vitro*, repeat-containing RNA can self-aggregate. Two structural motifs have been proposed to provide the interstrand interactions that drive aggregation: G-quartet (G4) structures and double-strand interactions with GG mismatches. Using *in vitro* transcribed RNA with pathologically relevant number of repeats, we were able to form gel-like aggregates suitable for investingation using fast MAS NMR spectroscopy. This approach enabled us to characterize the dominant interstrand interactions within the RNA gels. Both Watson–Crick and Hoogsteen base pairs were identified in RNA gels formed by RNA with 48 GGGGCC repeats. Their relative abundance shifted upon reconstitution in the presence of different divalent cations or nuclear extracts, underscoring the dynamic equilibrium between G-quadruplex and duplex interactions in GGGGCC RNA aggregation.

## Introduction

Expansion of GGGGCC repeat sequences within the *C9orf72* gene is associated with neurodegenerative diseases, such as amyotrophic lateral sclerosis (ALS) and frontotemporal dementia (FTD) ^1–4^. Repeat expansions with hundreds of repeats have been observed in some patients with FTD/ALS, whereas unaffected individuals have fewer than 24 repeats and most commonly only two repeats^1^. Disease mechanism of the mutation is quite complex and involves haploinsufficiency, RNA toxicity and/or toxicity of translated dipeptide repeats^5–9^. In the motor neurons of FTD/ALS patients and in model cell lines,GGGGCC RNA repeats have been shown to form nuclear foci, which have paraspeckle like properties^1,10,11^. It seems that in model cell lines, the threshold for foci formation is between 16 to 38 repeats^9,12^. In these studies, higher number of repeats induced foci that were larger and more numerous or present in a larger population of cells^9,12^. The threshold for repeat RNA aggregation is lower under *in vitro* conditions and as low as 5 repeats formed aggregates under *in vitro* conditions while extending repeats to 10 or more changed their appearance and ease of aggregation^9,12,13^.

The *in vitro* aggregation of RNA repeats raises the question of which structural motifs could be supporting the intermolecular association of RNA. The structure of GGGGCC repeats, both in DNA and RNA form, has been intensively studied in recent years. A wide variety of structures of GGGGCC repeats have been determined. G-quartets (G4) have been observed to fold from a single strand or assemble as a tetramer of GGGGCC oligonucleotides^6,14–21^. Hairpins and double-stranded structures with internal loops were also observed^15,22,23^. The double-stranded structures can form in two ways: They contain internal loops with either single GG mismatches or double GG mismatches^24–26^. All these structural motifs can occur both intramolecularly and intermolecularly. For long RNAs with a large number of interacting repeats, such interactions lead to phase separation and sol-gel transition^27^. This process can be even more complex *in vivo* due to additional interactions between RNA and protein binders that are multivalent ^9,15,16,28–30^.

The number of repeats is an important factor when studying the large-scale organization of repeat RNA^1,9^. Biophysical techniques such as CD, NMR spectroscopy, and X-ray crystallography have been successfully used to study RNA oligonucleotides with up to 8 GGGGCC repeats^6,16,17,23,31^. Obtaining structural information on longer RNAs, ideally with more than the pathological threshold of 24 repeats in length, could inform us on how multivalency affects structure.

We show that a technique called NMR spectroscopy under fast Magic Angle Spinning (fast MAS NMR) can be used to obtain structural information on GGGGCC repeat RNAs of pathologically relevant lenghts. Unique characteristics of MAS NMR, either conventional or with fast spinning rates, allow us to obtain structural information on biomolecules regardless of their size. This means that information can be obtained for samples in “solid” phases such as fibrils, microcrystals or sedimented samples^32–36^. MAS NMR has been previously used to study (CUG)_97_ repeat RNA^37,38^. Faster spinning rates during MAS NMR have the advantage of allowing ^1^H-detected experiments and Fast NMR MAS has been used to study a variety RNA structures^36,39–42^.

We used GGGGCC repeat RNA with 48 repeats (48xG4C2) to form aggregates that resemble gels and are suitable for very fast MAS NMR spectroscopy. Both Hoogsteen and Watson–Crick base pairs were present in these gels. The balance between the two shifted depending on the divalent cation added to induce aggregation or upon addition of nuclear extracts.

## Results

### Divalent ions trigger 48xG4C2 aggregation

Template DNA vectors for repeat RNA transcription were prepared as previously described^9^. The 48xG4C2 construct contained 48 repeats of GGGGCC (Figure 1A), preceded by a 25-base-pair flanking sequence containing all four nucleotides and ending with “GGG” at the 3′ end. The transcribed and purified 48xG4C2 formed gels upon addition of the divalent ions (Figure 1). Shorter RNAs with 8 repeats or 24 GGGGCC repeats formed smaller gels under the same conditions (Supplementary Fig. 1). Notably, both 24xG4C2 and 48xG4C2 formed multimeric species already during transcription, as evidenced by multiple bands and incomplete denaturation on TBE-urea PAGE, including smeared migration and RNA retained in the gel pockets (Supplementary Fig. 1, Supplementary Fig. 2). These observations indicate that the multimers are highly stable and resist full denaturation even in 7 M urea.

**Figure 1:**
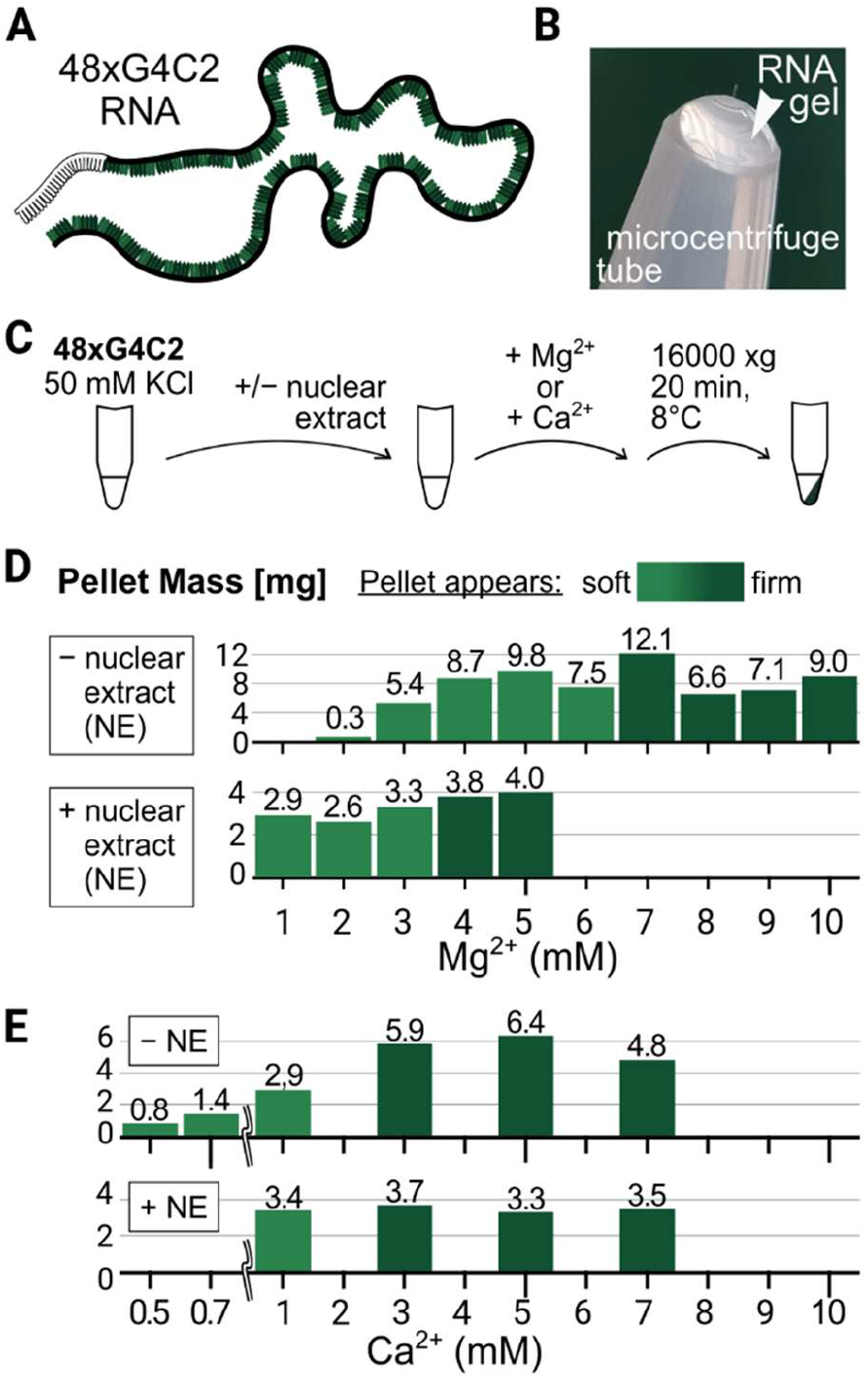
Identifying divalent-ion thresholds for reproducibly obtaining sufficient RNA-gel material for MAS NMR. A) Schematic of the transcribed 48xG4C2 construct; GGGGCC repeats are colored green. B) A representative photo of an inverted Eppendorf tube showing the 48xG4C2 gel. C) Gels were prepared by adding Mg^2+^ or Ca^2+^ to purified RNA and pelleting the material in a tabletop centrifuge. Panels D and E) Pellet properties as a function of ion concentration. Higher concentrations of divalent ions yielded larger pellets that appear firmer and less fluid. Pellet masses are given in milligrams above the bar. Shades of green indicate the apparent firmness of each pellet as assessed by visual inspection. Because half the amount of RNA was used for gel formation with nuclear extracts, the –NE and +NE panels are not directly comparable; the aim was to determine the ion concentration threshold rather than to compare pellet sizes across conditions.

Addition of divalent ions triggered gel formation by 48xG4C2 RNA (Figure 1B), using the buffer and centrifugation conditions outlined in Figure 1C. All gel formation experiments were performed in the presence of 50 mM KCl, a concentration chosen to approach intracellular ionic conditions and to provide conditions for G-quadruples folding. In the presence of Mg^2+^ (Figure 1D), at least 3 mM Mg^2+^ was required to induce substantial aggregation, and higher concentrations of MgCl_2_ yielded gels that were larger and firmer. Although these pellets likely consisted of a highly hydrated RNA gel containing a substantial amount of water, the RNA:water (w/w) ratio was not quantified. Compared to magnesium, calcium ions were more efficient at promoting aggregation, with gel formation observed at concentrations as low as 5 mM CaCl_2_ (Figure 1E), and higher concentrations (>3 mM) caused significant changes in size and appearance.

The addition of nuclear extract (HeLaScribe® Nuclear Extract, Promega, E3092) to 48xG4C2 prior to the addition of MgCl_2_ lowered the concentration threshold required for aggregation of 48xG4C2 (Figure 1D). This effect was not observed when CaCl_2_ was added to 48xG4C2 pre-mixed with nuclear extracts (Figure 1E). Since our goal was to determine divalent-ion thresholds for reproducibly obtaining sufficient RNA-gel material for MAS NMR, the total RNA amounts were not matched to buffer-only conditions, and the resulting gel masses are therefore not directly comparable between divalent-ion series. In addition, both the presence of nuclear extract and the choice of divalent ion may have affected the water content of the pellets, making direct mass comparisons difficult. Finally, the nuclear extract contained insoluble material that remained in these tests but was removed for MAS NMR gels.

### NMR spectroscopy of model RNA repeats

The GGGGCC repeats have the ability to form various structures, such as double-stranded RNA through Watson–Crick base-pairing, hairpins, and also G4s through Hoogsteen base pairing. To obtain representative NMR spectra that could later be compared with those of 48xG4C2, we designed two model oligonucleotides: a G4 and a double-stranded RNA with an internal loop (Figure 2A, 2B). The G4 forming sequence was (GGGGCC)_3_GGGG, allowing for a monomolecular G4 formation. We have screened RNA sequences that would form G4 through tetramerization of RNA strands as well (Supplementary figure 4). The double-strand sequence CCGGGGCCGG was designed to form a symmetric homodimer of with a double GG mismatch in the middle. After synthesis, the oligonucleotides were refolded under conditions that resulted in the most homogenous specturm. To obtain the G4 fold, we diluted the oligonucleotide to 10 µM, heated it to 95 °C, and then slowly annealed it to room temperature. The double-stranded homodimer was annealed at high concentration in the presence of 50 mM NaCl.

**Figure 2:**
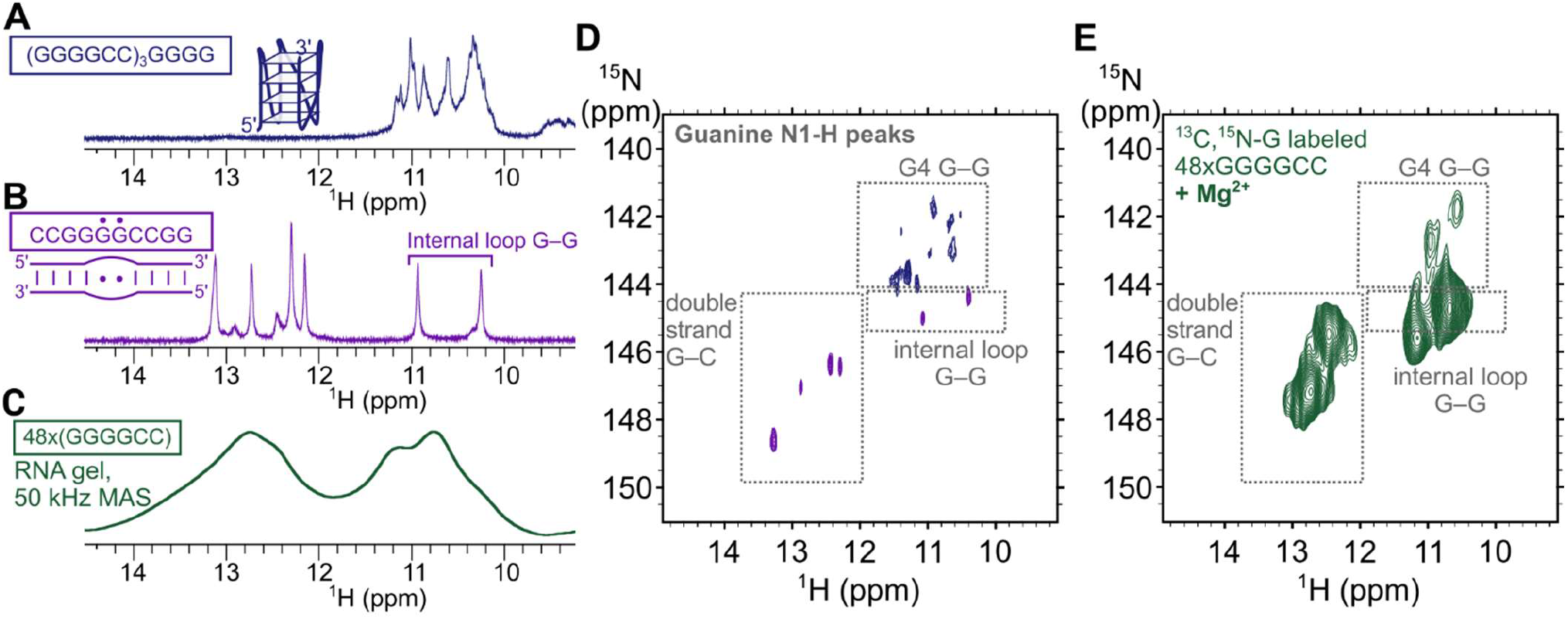
Solution-state NMR spectra and fast MAS NMR spectra of model RNA oligonucleotides and 48xG4C2 RNA gels. 1D ^1^H spectrum of an oligonucleotide folding into a G-quadruplex with chemical shifts characteristic of G4 Hoogsteen base pairs. B) 1D ^1^H spectrum of an oligonucleotide that dimerizes into an antiparallel duplex with an GG internal loop. In the schematic, vertical lines indicate Watson-Crick base pairing, and dots indicate a GG mismatch segment. The GG mismatch can form non-canonical G–G base pairs. C) Imino selective 1D ^1^H MAS NMR spectrum of 48xG4C2 RNA gel, showing signals from both G4 and duplex regions. D) SOFAST HMQC ^1^H–^15^N correlation spectrum of the double-stranded RNA oligonucleotide (violet) and G-quadruplex oligonucleotide (blue). Dashed boxes outline the chemical-shift regions characteristic of distinct G– G and G–C base-pairing motifs. E) MAS NMR hNH spectrum of 48xG4C2 RNA gel formed in the presence of 10 mM MgCl_2_.

We recorded solution-state ^1^H NMR spectra of these two model RNA oligonucleotides (Figure 2A, 2B). The G4 fold gave rise to signals in the region between 10.4 and 11.5 ppm (Figure 2A). These chemical shifts correspond to imino protons of G-tetrads formed through Hoogsteen base pairs^14,18,43^. Regardless of whether the G4 is parallel or antiparallel, monomolecular or multistranded, the imino proton involved in the G–G base pair gives a characteristic signal between 10.5 and 12.0 ppm (Supplementary Figure 4)^17,21,44^.

The double-stranded dimer gave rise to a series of signals in the region between 12.0 and 13.4 ppm (Figure 2B). This is typical of G–C Watson–Crick base pairs in a double-stranded DNA/RNA^45^. Two additional signals appeared at around 10.4 and 11.9 ppm, indicating G–G pairing inside an internal loop as previously observed in the literature^24–26^.

While the internal-loop G–G pairs overlap with the G4 in the imino-proton region, the two model oligonucleotides exhibited distinct ^15^N chemical shifts (Figure 2D). All G4-related cross-peaks appeared below δN 144.1 ppm. Characteristic ^15^N chemical shifts of G4 have been reported previously^46–48^.

In summary, the spectra of the model oligonucleotides showed that double-stranded and G4 structures each gave rise to well resolved signals with distinct chemical shifts. By leveraging the additional ^15^N dimension in 2D ^1^H–^15^N correlation spectra, the GG-mismatch and the G4 cross-peaks could be distinguished.

### Obtaining fast MAS spectra of RNA gels

For solid-state NMR spectroscopy measurements, we prepared 48xG4C2 RNA by transcription in the presence of ^13^C,^15^N-GTP, resulting in 48xG4C2 containing ^13^C,^15^N-isotopically labeled guanines at all positions. This RNA was aggregated into gels in the presence of 10 mM MgCl_2_. Between 5 and 10 mg of RNA pellet were packed into 1.3 mm MAS NMR rotors (3 µl volume) to account for losses and gel compaction. We recorded an imino selective 1D-^1^H MAS NMR spectrum of 48xG4C2 RNA gels. As expected, the highly heterogeneous nature of the 48xG4C2 gels combined with spinning rate of 50 kHz gave rise to broad cross-peaks (Figure 2C). We could not determine with certainty whether G4 formation occurred in the 48xG4C2 gels. To obtain better resolution, we recorded 2D ^1^H–^15^N correlation MAS NMR spectra of 48xG4C2 gels (Figure 2E). The additional ^15^N dimension allowed us to more clearly map the fingerprints of model oligonucleotides to the spectrum of 48xG4C2 which overlapped better with the peaks of the double-stranded model oligonucleotide. A weak cross-peak density was observed in the region corresponding to the G4 fold (a chemical shift below δ_N_ 144 ppm). Although the cross-peaks corresponding to the G4 fold were weaker than others, the data suggest coexistence of Watson–Crick base-paired guanines and Hoogsteen base-paired guanines (Figure 2E).

Given that several RNA structural motifs are, in principle, capable of supporting multimerization^49^, both detected interaction types could contribute to gel formation (Figure 3A). One possible model is that both RNA duplexes and parallel G4 bring together strands of different RNA molecules. Additionally, the observed peaks could reflect contributions from hairpin structures or monomolecular G4 to some extent, since based on chemical shifts do not allow us to distinguish between intramolecular interactions within a single RNA strand and intermolecular interactions in trans between distinct RNA molecules. However, because dynamics influence signal strength, only structures that are part of rigid, large complexes are likely to generate detectable signals under MAS NMR conditions, meaning that hairpins embedded within flexible single strands are less likely to significantly contribute to the observed peaks.

**Figure 3:**
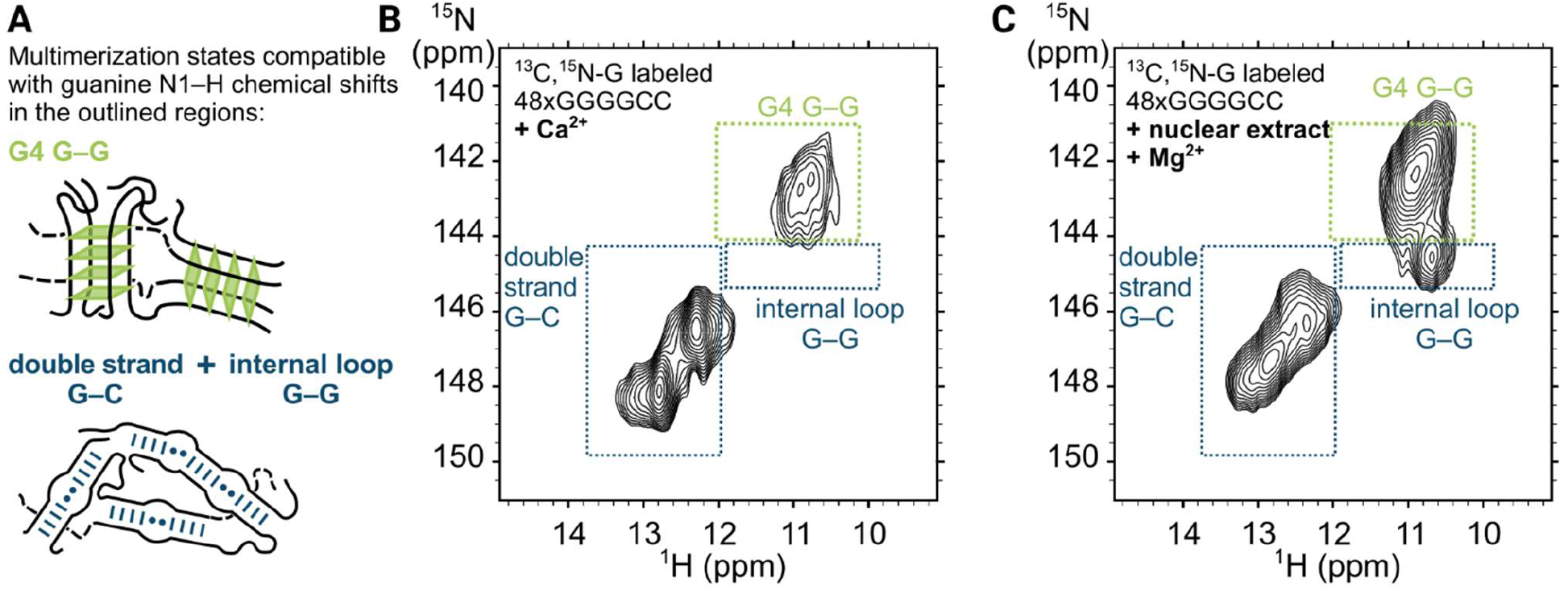
2D ^1^H–^15^N correlation spectra of aggregated 48xG4C2 RNA highlight differences in regions corresponding to G4 quartets formed via Hoogsteen base pairs. A) Schematics of two potential multimerization modes. Each mode has a distinct base-pairing arrangement resulting in characteristic guanine N1-H chemical shifts. B) MAS NMR hNH spectrum of 48xG4C2 RNA gel formed in the presence of 10 mM CaCl_2_. Rectangles outline chemical-shift regions corresponding to the multimerization states shown in panel A. C) MAS NMR hNH spectrum of 48xG4C2 RNA gel formed in the presence of nuclear extracts and 10 mM MgCl_2_. The corresponding motif-specific regions are highlighted for comparison with panel A.

### The effect of CaCl_2_ on the spectra of RNA gels

We formed 48xG4C2 RNA gels in the presence of CaCl_2_ and measured 2D ^1^H–^15^N correlation MAS NMR spectra (Figure 3B). The spectrum was similar to that of 48xG4C2 gels formed in the presence of MgCl_2_, but with more intense and prominent cross-peaks corresponding to the G4 fold (chemical shift below δ_N_ 144 ppm). The peak density centered at around δ_H_ 11 ppm fit-in well into the outlined G4 region in terms of nitrogen chemical shifts. Based on this observation, it is possible that the gels formed by presence of CaCl_2_ have a greater proportion of G4 stacking interactions than those formed in the presence of MgCl_2_. However, the double-strand related G-C interactions remained the dominant structural feature based on dominating peak density in the downfield region (Figure 3B, between δ_H_ 12 and 13 ppm and δ_N_ 145.5 and 149.0 ppm). It is also worth noting that the absence of peaks for internal loop G–G base pairs does not imply that internal loops are absent, as changes in geometry, heterogeneity, or dynamics could prevent their signal strength.

### The effect of nuclear extracts on RNA gels

Next, we wanted to see if components from the cell nuclei could alter the conformation of the 48xG4C2 RNA. The insoluble fraction of the nuclear extracts was removed by centrifugation (10 min, 1,000 ×g, 15 °C), and the clarified nuclear extracts were added at a volume ratio of 1:1 to ^13^C,^15^N-guanine labeled 48xG4C2 (Figure 1C). After a short incubation period, we induced gel formation by adding 10 mM MgCl_2_. We then performed two washing steps (resuspension and pelleting) to remove the non-bound nuclear components. We packed the resulting 48xG4C2 gel, formed in the presence of nuclear extract, into a 1.3 mm MAS NMR rotor and acquired ^13^C-^15^N correlation spectra (Figure 3C). The cross-peaks mapped well to both the G4 and double-strand model oligonucleotides, without a clear preference for either of them (Figure 3C). Western blotting and SDS-PAGE analysis confirmed that proteins remained bound to 48xG4C2 gel after washes (Supplementary Fig. 3).

## Conclusion

The aim of this study was to assess the suitability of fast MAS NMR for investigating RNA with a pathologically relevant number of GGGGCC repeats. We successfully identified conditions for preparing gel-like aggregates from RNA that would be suitable for analysis using solid-state NMR spectroscopy. We established an analytical approach, though still qualitative in nature, that can give us information on structure of repeat RNA in gel-like aggregates. Using fast MAS NMR we were able to observe imino secondary shifts of guanines, informing us on Watson-Crick and Hoogsteen bond formation.

We prepared ^13^C,^15^N-labeled 48xG4C2 RNA, which formed gel-like aggregates upon the addition of 10 mM Mg^2+^ or 10 mM Ca^2+^ ions. While these conformations are not physiological, the structural data provided interesting insights. In the presence of 10 mM Mg^2+^, the double-stranded conformation was dominant. When Mg^2+^ was replaced by Ca^2+^, the absence of Mg^2+^ possibly reduced the RNA duplex stability, leading to a greater fraction of G4 structures. Gels formed in the presence of nuclear extracts contained both conformations, the double-strand and G4. Taken together, the data indicate that duplex and parallel G4 interactions may simultaneously contribute to gel formation.

Access to both ^1^H and ^15^N chemical shifts by using fast MAS NMR allowed us to distinguish the guanine bonding geometries present in 48×G4C2 gels and, by extension, the multimerization interactions they support. Although the intrinsic structural heterogeneity of GGGGCC repeats prevents unambiguous identification of all species, the resulting restraints are well suited for validating existing multimerization models against data from RNA with pathologically relevant repeat lengths or characterizing ion effects on multivalent RNA-RNA interactions.

## Materials and Methods

### RNA transcription and purification

The pcDNA3 plasmids containing G4C2 repeats in multiples of 8, 24, and 48 were previously constructed and described elswhere^9,10^. Transcription templates were generated by linearizing the plasmids using the BamHI restriction enzyme (NEB, cat. no. R3136L), rather than by PCR amplification. Linearized products were purified and concentrated Zymo DNA Clean and Concentrator - 500.

RNA transcribed from 48xG4C2 plasmid contained a starting sequence of GAGACCCAAGCTTGTGTGGTTTA, followed by 48 G4C2 repeats, and terminated with GGG at the 3’ end.

Following small scale optimizations (100 µl – 1 ml) to ensure that variations in buffer and ion composition did not significantly affect RNA yield we selected a potassium-containing transcription buffer (20 mM HEPES pH 7.9, 100 mM KCl, 3 mM MgCl_2_, 1 mM spermidine, 10 mM DTT, 0.05 mM EDTA, YIPP at 1:150000 dilution, 500 µM of each rNTP).

For solid-state NMR samples, rGTP was replaced with ^13^C,^15^N-rGTP. Transcription reactions were done in 10 ml final volume, with 500 µg of linearized template, 333 µl (16’500 units) of T7 RNA polymerase (NEB, cat. M0251L). Transcription mixture was incubated at 37°C for 4 hours. Turbo DNase (Invitrogen cat no. AM2238) was added (63 µl, 126 units) to digest the DNA template.

The transcribed RNA was purified using dialysis followed by phenol-chloroform extraction with the aim to remove excess nucleotides and proteins, including ^13^C,^15^N-rGTP that might contribute the signal to NMR spectra. Dialisys was done against dialysis buffer (20 mM HEPES pH 7.9, 100 mM KCl, 3 mM MgCl2, 2 mM DTT) first for 2 hours with 500 ml and overnight with 500 ml buffer. Dialysis was done in Slide-A-Lyzer cassettes with a 15 ml volume and MWCO 3.5 kDa (ThermoFisher Scientific, cat no. A52967). Subsequently RNA was purified via phenol–chloroform extraction. The resulting pellet was reddisolved in 300 µl water giving final concentrations in the range between 12 µg/µl and 15 µl/µl. Total RNA yield from a typical 10 ml transcription reaction was between 3.5 mg and 4.5 mg.

### RNA gel formation in the presence of KCl and divalent ions

Test precipitations were performed in microcentrifuge tubes at a final volume of 30 µl containing 20 mM HEPES, 100 mM KCl, and varying concentrations of MgCl_2_ or CaCl_2_. The final concentration of RNA was maintained at 6 µg/µl (total volume: 30 µl, total RNA: 180 µg). After a 10-minute incubation at 4 °C, the samples were centrifuged at 16,000 ×g at 4 °C to pellet the formed RNA gel. After removing the supernatant, the resulting pellets were weighed to determine their mass. The water content in the formed RNA gels was not quantified. Samples of pure gelified RNA used for MAS NMR spectroscopy were prepared under the same conditions.

The HeLa Scribe Nuclear Extract (Promega, cat. E3110, now discontinued) was used to examine the effects of nuclear factors in gel formation. Nuclear extract buffer composition was 40 mM Tris pH 8.0, 100 mM KCl, 0.2 mM EDTA, 0.5 mM PMSF, 0.5 mM DTT, 25 % glycerol.

### RNA gel formation in the presence of nuclear extracts for NMR spectroscopy

Before gel formation, nuclear extract was first clarified by centrifugation at 1,000 ×g for 10 minutes at 15 °C. This removed insoluble material that would otherwise end up in RNA gels. Next, 28 µl of ^13^C,^15^N-rGTP labeled RNA (containing 100 mM KCl and 1 mM DTT) was pre-incubated with 30 µl of clarified nuclear extract for 10 minutes at room temperature. The mixture of RNA and nuclear extract was then added to a new tube containing 2 µl of 300 mM MgCl_2_ and vigorously mixed. This resulted in a final reaction volume of 60 µl with the final buffer composition of 20 mM Tris pH 8.0, 90 mM KCl, 0.1 mM EDTA, 0.25 mM PMSF, 0.75 mM DTT, 12.5% glycerol, and 10 mM MgCl_2_. To remove buffer components such as Tris, that could be detected in NMR spectra, resulting gel was washed twice with 250 µl wash solution (20 mM HEPES, 100 mM KCl, 10 mM MgCl_2_, 1 mM DTT, 10% ^13^C-depleted glycerol).

### Synthesis and refolding of model oligonucleotides

Oligonucleotides were synthesized on a DNA/RNA synthesizer H8 (K&A Laborgeraete GbR, Germany) using standard solid-phase phosphoramidite chemistry in DMT-on mode. Oligonucleotides were cleaved from the solid support and fully deprotected using an aqueous ammonia/methylamine (AMA) solution (1:1 v/v) at 65 °C for 15 minutes. The RNA was lyophilized, and the TBS protecting groups were removed by dissolution in dimethyl sulfoxide (DMSO), followed by incubation with triethylamine (TEA) and triethylamine trihydrofluoride (TEA·3HF) at 65 °C for 15 minutes. The RNA oligonucleotides were then immediately purified using GlenPak RNA cartridges following the standard GlenPak procedure. Finally, the samples were lyophilized and resuspended in H_2_O (HPLC grade). The concentration of oligos was determined spectroscopically (OD_260_).

The oligonucleotides folding into a quadruplex structure with a sequence (GGGGCC)_3_GGGG was diluted to 10 µM in 10 mM KCl in H_2_O. After heating to 95 °C for 5 minutes in a water bath, the sample was left in the switched-off water bath overnight to cool to room temperature. The annealed oligonucleotide was concentrated using Amicon ultrafilters.

The oligonucleotide folding into the duplex with a sequence CCGGGGCCGG was refolded without dilution at concentrations greater than 100 µM in the presence of 50 mM NaCl and 0.5 mM MgCl_2_. After heating to 95 °C for 5 minutes, the sample was cooled to room temperature by removing the microcentrifuge tube from the water bath.

### Solution state NMR spectroscopy

Solution-state NMR titration experiments were performed using Bruker AVANCE NEO 600 MHz spectrometers equipped with cryogenic probes. 1D ^1^H NMR experiments were performed with a spectral width of 19 ppm, 512 scans and a relaxation delay of 1 sec. Water suppression was achieved using excitation sculpting. Samples for NMR were prepared at concentrations ranging between 0.1 mM and 0.8 mM RNA in 95% H_2_O/5% ^2^H_2_O supplemented with 10 mM potassium phosphate buffer (pH 7.0) and 10 mM KCl. The RNA samples were annealed at 90 °C for 5 minutes and slowly cooled overnight.

### Rotor packing and Solid state NMR spectroscopy

RNA gels were packed into 1.3 mm rotors (3 µl volume) by centrifugation (67,000 ×g, 16 hours, 10 °C). Before packing, a 10 mg RNA gel was resuspended in appropriate buffer (100 mM KCl, 10 mM MgCl2 for purified samples, and wash buffer for RNA with nuclear extracts). Gel resuspension was pelleted into the rotor with a specifically designed insertion tool (GiottoBiotech). NMR experiments were conducted on a 23.5 T Bruker spectrometer (ω_0_,H/2π = 950 MHz) equipped with a 1.3 mm HCN probe and an NEO console. During measurement, the rotor was spun at 50 kHz. Frictional heating was compensated with a gas flow of 1350 lph, at a stator temperature of 250 K, resulting in an estimated internal sample temperature at 35 °C. The 2D ^1^H–^15^N and ^1^H–^13^C correlation spectra were acquired using standard ^1^H-detected Cross-Polarisation (CP, linear ramp) based HSQC experiment (available in NMRlib 2.0 as av_hNH_cp_cp_miss.IBS).

The 48xG4C2 gel formed in the presence of CaCl_2_ was measured at the CRMN Lyon facility under slightly modified conditions (e.g., sample packing and spinning rate). RNA gels were packed into 1.3 mm rotors by centrifugation (150,000 ×g, 1.5 hours, 10 °C). The NMR experiments were conducted on a 23.5 T Bruker spectrometer (ω_0_,H/2π = 1,000 MHz) equipped with a 1.3 mm HCN probe. During measurement, the rotor was spun at 60 kHz. Frictional heating was compensated with a gas flow of 1000 lph, at a stator temperature of 240 K, resulting in an estimated internal sample temperature at 25 °C. The 2D ^1^H–^15^N and ^1^H–^13^C correlation spectra were acquired using standard ^1^H-detected Cross-Polarisation (CP)-based HSQC experiment.

### Immunoblotting

48xG4C2 samples were diluted with precipitation buffer lacking MgCl_2_ (20 mM HEPES pH 7.9, 100 mM KCl, 2 mM DTT, 5% glycerol). Dissolution of 48xG4C2 gels improved with the addition of 0.2 mM EDTA. Loading amounts were estimated based on the final dilution factor of the nuclear extract. Samples and the All Blue Prestained Protein Standard (Bio-Rad, 1610373) were loaded onto 12% stain-free gels (Bio-Rad, Hercules, California, USA) and transferred to nitrocellulose membrane using the semi-dry transfer method.

Blocking was performed for 1 hour at room temperature in blocking solution (5% skim milk in TBS with 0.05% Tween-20). Membrane was incubated overnight at 4°C with the primary anti-hnRNPH antibody (Proteintech, 67375-1-Ig) diluted 1:1000 in blocking solution. The next day, the membrane was washed three times for 10 min with TBST, then incubated with Peroxidase AffiniPure Goat Anti-Rabbit IgG (H+L) secondary antibody (111-035-144, Jackson ImmunoResearch) diluted 1:5000 in blocking solution. Clarity Max Western ECL Substrate (Bio-Rad, Hercules, California, USA) was used for signal detection and images were captured using GelDoc System and ImageLab software (Bio-Rad, Hercules, CA, USA).

## Additional author information

### Author contributions

Sara Zager prepared G4C2 RNA, synthesized RNA oligos, and conducted gelation assays. Nataša Medved synthesized RNA oligos. Mirko Cevec carried out solution-state NMR measurements. Urša Čerček prepared plasmids, performed western blots, and contributed to manuscript revisions. Boris Rogelj and Janez Plavec provided project guidance and feedback, as well as critical manuscript revisions. Jaka Kragelj designed the project goals, established the protocols for sample preparations, prepared G4C2 RNA and samples for MAS NMR, analyzed data, and authored the manuscript.

## Acknowledgments

We would like to thank Tanguy Le Marchand from CRMN Lyon and Alicia Vallet from IBS Grenoble for performing MAS NMR spectroscopy measurements. We would like to thank the members of the Jernej Ule lab, particularly Tajda Klobučar, for their assistance with analyzing RNA sample quality and performing analytical RNA purifications. Additionally, we thank Marjetka Podobnik, Head of the Department of Molecular Biology and Nanobiotechnology, for providing access to microbiology equipment, and Tanja Peric for her help with the cultures required for plasmid purification.

