## Supplemental Figures 1 to 4 for "Fast MAS NMR Spectroscopy Can Identify G-Quartets and Double-Stranded Structures in Aggregates Formed by GGGGCC RNA Repeats"

Supplementary info for:

### RNA containing GGGGCC repeats forms foci mostly through double strand interactions and through G-quadruplet stacking as well.

Supplementary Figure 1

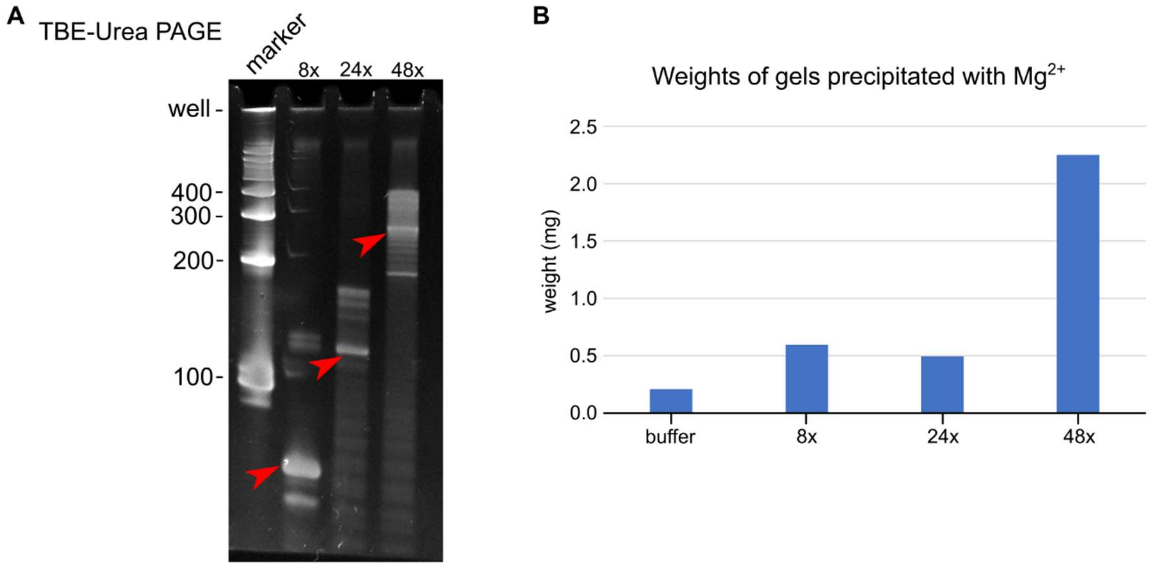


*Supplementary Figure 1: Transcribed GGGGCC repeat RNA of different lengths does not form gels of equal size (weight) upon precipitation with 10 mM $MgCl_2$. A) TBA Urea PAGE analysis of transcribed 8xG4C2, 24xG4C2, and 48xG4C2. Due to multimerization 48xG4C2 displays several higher molecular weight bands on the gel, as does 24xG4C2, showing that heating in the presence of Urea does not completely denature the multimeric structures. Bands of lower molecular weight than expected are presumably due to partial folding and possibly early termination, which represents a minor fraction of total RNA. B) Weight of the microcentrifuge tubes after gel formation and after the supernatant was aspirated from the microcentrifuge tubes.*

Supplementary Figure 2

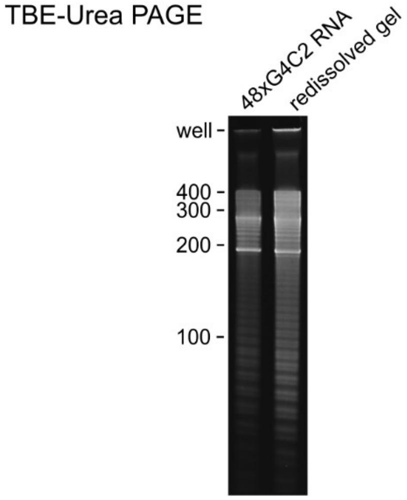


*Supplementary Figure 2: Confirmation that the gels formed with 48xG4C2 RNA indeed contain RNA. Samples were analyzed on were analyzed on TBE Urea PAGE gel. Lane 1) The 48xG4C2 RNA in soluble form before precipitation. Lane 2) The 48xG4C2 gel redissolved in buffer without magnesium and loaded on the gel. Note the incompletely denatured 48xG4C2, that remains in the gel pocket and does not migrate on the TBE-Urea PAGE.*

Supplementary Figure 3

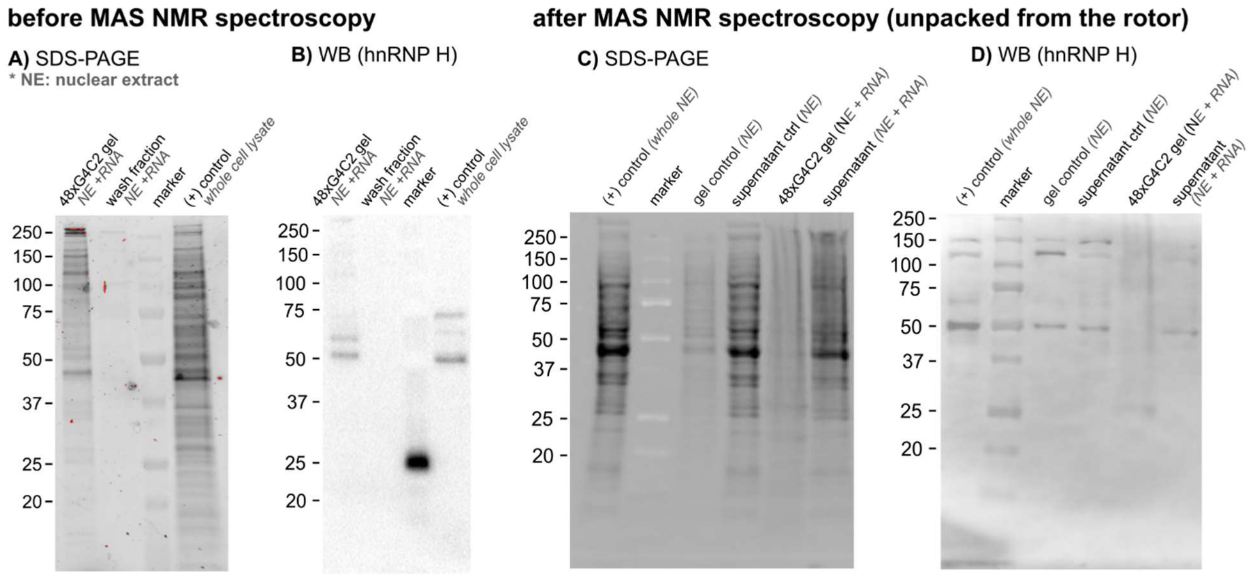


*Supplementary Figure 3: Some proteins from the nuclear extracts get trapped in the 48xG4C2 gel including the hnRNP H as confirmed by western blot. A, B) Examination of whether nuclear extract proteins remain bound to the 48xG4C2 gel remained bound after three washes of buffer. C, D) Assessment of excess stoichiometry of nuclear extract. We examined whether nuclear extract proteins are in excess stoichiometry and therefore remain in supernatant. The 48xG4C2 gel did not completely dissolve after packing into the rotor and after the MAS NMR spectroscopy measurements, possibly due to insufficient dilution leaving residual $Mg^{2+}$ or perhaps due to something specific to the MAS NMR spectroscopy conditions.*

Supplementary Figure 4

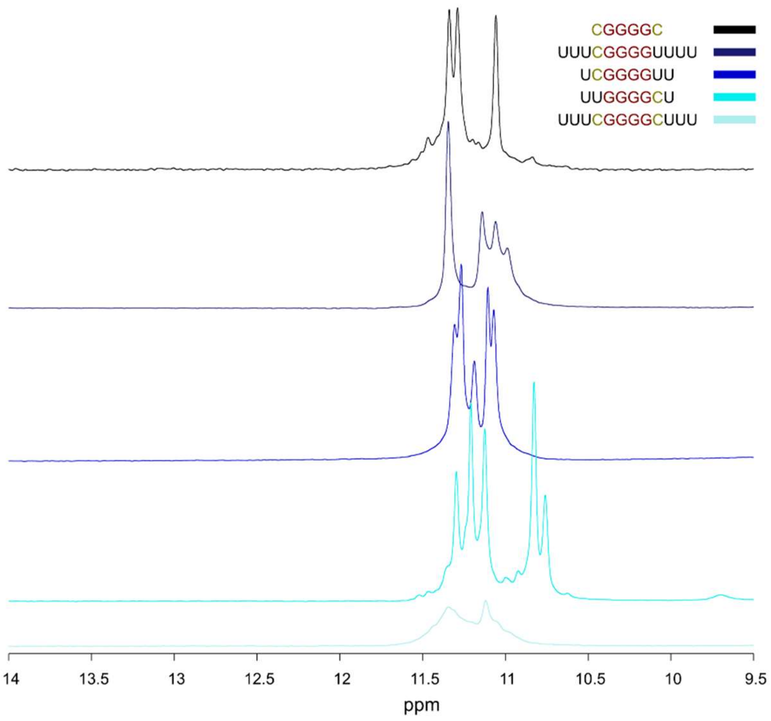


*Supplementary Figure 4: $^{1}$H NMR spectra of screened GGGG RNA sequence variants to determine the best flanking sequence for G4 formed through tetramerization. These oligonucleotides are expected to form parallel G4 through tetramerization. These oligonucleotides were annealed in the presence of 50 mM KCl.*
